## Supplemental Information for "Loop-extruders alter bacterial chromosome topology to direct entropic forces for segregation"

November 28, 2023

### 1 Blob argument for cost of strand overlap

In the main text we considered the entropically preferred states of the largest ring and the linear segment independently when arguing that fork-segregated states are entropically preferred. Here, we consider excluded volume interactions between these components, and show that fork segregation can still be entropically favorable at intermediate replication stages. In brief, we will show that since fork-segregated states contain regions where only *three* polymer chains overlap along the long axis of the cell (Main Fig. 1A), whereas sister strand-segregated states contain regions where *four* chains overlap (Main Fig. 1B), fork-segregated states can also minimize the entropic cost due to chain overlap.

To show that fork segregation can also reduce the entropic cost of strand overlap, we first provide a blob argument for the preferred configuration of a half-replicated chromosome ($R \approx N/2$) confined to a tube. Following [1], we assume that the free energy cost of confinement scales with the number of blobs the polymer splits into, and that when $n$ polymer strands are parallel in a tube of diameter $d$, they can be modeled as being confined into effective tubes of diameter $d/\sqrt{n}$. In this picture, the number of blobs per unit length is given by

$$\mathcal{B}(n) = \frac{n\sqrt{n}}{d}. \tag{1}$$

This immediately shows that it is entropically costly to have more polymer strands in parallel along the tube, since the average blob size is smaller.

We now consider the cost of chain overlap of a segregated state and a fork-segregated state without loop-extruders. In the segregated state (Sup. Fig. 1A), one replicated strand is doubled along the long axis and overlaps with the unreplicated region, creating one region of length $fL$ with $n = 4$ parallel strands, and another of length $(1 - f)L$ with $n = 2$ parallel strands.

To find $f$, the fraction of the length occupied by four parallel strands, we use that the replicated strands are of the same length, so that the same number of monomers must fit into each strand. This implies that

$$f = \frac{1}{1 + \sqrt{2}^{1-1/\nu}} \approx 0.56. \tag{2}$$

This result is intuitive: In the region with more parallel strands, the same amount of monomers are confined to a narrower tube, and hence the region is more extended, i.e. $f > 0.5$.

We can use this result to estimate the total number of blobs for the segregated configuration:

$$B_{\text{segregated}} = \mathcal{B}(2)(1 - f)L + \mathcal{B}(4)fL \approx 5.7 \times L/d. \tag{3}$$

This is larger than in the fork-segregated state, where all three strands are in parallel (Sup. Fig. 1B):

$$B_{\text{fork-segregated}} = \mathcal{B}(3)L \approx 5.2 \times L/d. \tag{4}$$

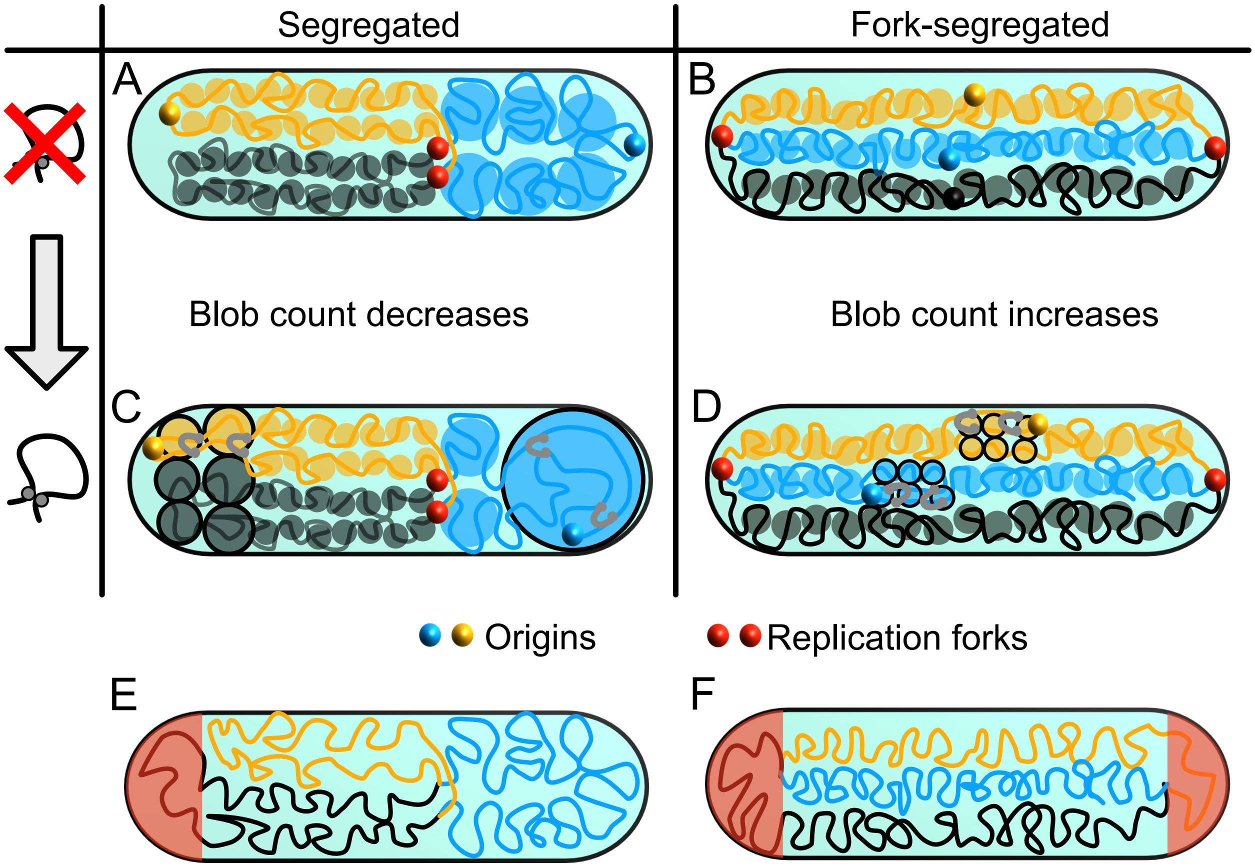


Figure 1: **Polymer blob sketches for $R = N/2$** **A** A segregated chromosome configuration. A fraction $f$ of the volume is occupied by four chromosome strands in parallel, and a fraction $(1 - f)$ is occupied by two strands in parallel. Since the replicated strands are of equal length, it is easily seen that $f > 0.5$. **B** In the fork-segregated state, three chromosomal strands are extended in parallel. **C** When loop-extruders are added, the origin-proximal regions are linearized, and start to behave as single polymer strands. This means that in the segregated state, the two ends of the subvolume now each have three or one parallel polymer strands. **D** In the fork-segregated state, the linearized segments have to bend in either direction, meaning that a region with four parallel polymer strands is created. **E** Possible overhang (within red-shaded region) for a sister-segregated state. An overhang at the pole with four strands can reduce the cost of strand overlap. Note that this is the only possible overhang; due to the topology of the chromosome, displacing any other strand will not reduce the number of overlapping strands in the configuration. **F** Possible overhangs (within red-shaded regions) for a fork-segregated state. At both replication forks, one of the three chromomal strands could cause an overhang that reduces the entropic cost of strand overlap.

Although the resulting free energy difference due to overlap ($k_{\mathrm{B}}T$ per blob) is relatively small, we note that we have neglected the bending of the unreplicated strands in the segregated state, which is expected to further increase the free energy cost for the segregated state. Constraints near the replication forks are neglected for both cases.

For simplicity, we have neglected the possibility of "overhangs" (where one strand extends beyond the region of overlap [2, 3, 4, 5]) in both the fork-segregated and the sister-segregated state. An overhang can reduce the entropic cost of overlap by a constant factor independent of the tube diameter or chain length [2, 3, 4]. Since fork-segregated states could have two overhangs (one at each replication fork, Sup. Fig. 1F) whereas sister-segregated states could have only one overhang (at the pole with four strands Sup. Fig. 1E), we do not expect overhangs to change the entropically preferred global orientation.

A similar blob analysis could be made for other replication stages, but the unequal lengths of the strands would have to be taken into account, as done in [5] for two linear polymers. Qualitatively, however, a similar argument should hold: a sister-segregated state would only be preferable compared to a fork-segregated one if the fraction of the tube occupied by four parallel strands, $f$, would be small enough. However, since the replicated strands are of equal length, the strand confined to a narrower tube should be more extended, hence $f > 0.5$. Detailed Monte Carlo simulations–as performed in [4, 5]–to better assess the free energy landscape for partially replicated bacterial chromosome fall outside the scope of our current work.

### 2 Blob number changes with loop-extrusion

We now consider what happens to the number of confinement blobs when loop-extruders are loaded at the origins of replication. Suppose the loop-extruders linearize a region of genomic length $\lambda$ near the origins, by effectively zipping together the two chromosomal arms around the origin. This linearized region will thus be referred to as a doubled-up strand. We will first assume that a doubled-up strand behaves effectively as a single polymer. In the segregated configuration, linearization of the *ori*-proximal regions near the cell poles *reduces* the number of parallel blobs (Sup. Fig. 1C). This means that loop-extruders make the segregated configuration more favorable. In the fork-segregated case, by contrast, the origins of replication are located mid-strand, so that the linearized regions will have to bend in either direction (Sup. Fig. 1D). This bending causes more strand overlap, and hence *increases* the number of blobs. The fork-segregated configuration hence becomes less favorable.

It cannot be assumed that the free energy cost per blob is the same for single chromosomal strands and doubled-up strands; since the doubled-up strand has more degrees of freedom, its free energy cost per blob is expected to be be higher. The overlap of doubled-up strands is hence entropically more costly, which further disfavors fork-segregated states compared to sister-segregated ones when loop-extruders are present.

### 3 Simulation parameters

For 3D polymer simulations, various parameters, such as the volume of confinement, need to be defined. Since our goal is to explore the possible roles of various segregation mechanisms, rather than to model a specific bacterial species, we base our parameter choices on *Caulobacter crescentus* and *Bacillus subtilis*, since a lot is known about these two species. However, the sensitivity of our main results to these chosen parameters is explored thoroughly.

*C. crescentus* is a model organism where the ParAB*S* system's role in chromosome segregation has been extensively studied. On the other hand, since previous simulation works have focused on modeling loop-extrusion in *B. subtilis*, we used these previously found simulation parameters for the loop-extruder number, processivity, and by-passing rate [6, 7]. Where relevant, we varied these parameters to test the robustness of our results (Sections 10, 11).

The monomer length corresponding to 10 kb was constrained using previously published microscopy data for *C. crescentus* [8]. In addition, we needed to constrain the replication fork and loop-extrusion speeds, as well as the cell size and *ori* positions at different replication stages.

#### 3.1 Cell size

Most bacteria grow exponentially in volume. The radius of *C. crescentus* cells remains fairly constant over the cell cycle [9], and we hence assume that only the length of the cells increases in time. The radius of the confinement is hence kept at a constant 0.63µm [8]. For the confinement length, we assume exponential growth

$$L(t) = L_0 e^{r_{\mathrm{L}} t}, \tag{5}$$

using $L_0 \approx$ 2.3 µm [8] and $r_{\mathrm{L}} \approx 5.5 \times 10^{-3}\ \mathrm{min}^{-1}$, so that the cell size doubles in 126 minutes [10, 11].

#### 3.2 *ori* separations

To define origin-pulling forces for the simulations, we required knowledge of how quickly replicated *ori*s are separated. The distance between the origins in *C. crescentus*, $d_{\mathrm{ori}}$, initially increases very rapidly, after which it increases gradually [12, 13, 14]. We fitted experimental data [15] with a function where the origins first separate with a starting speed of $v_0 \approx 330\ \mathrm{nm\ min}^{-1}$ and constant deceleration of approximately $a = -30\ \mathrm{nm\ min}^{-2}$ for 10 minutes, and then a constant velocity of roughly $v_f = 20\ \mathrm{nm\ min}^{-1}$(Fig.2B).

Assuming that one *ori* remains tethered in place, we used a tethering force to tie the stationary *ori* to a height of $z = 343$ µm [8]. The second *ori* was tethered to a height of $343 + d_{\mathrm{ori}}$, where $d_{\mathrm{ori}}$ was calculated by

$$d_{\mathrm{ori}}(t) = \begin{cases} v_0 t + \frac{at^2}{2}; t \leq 10\mathrm{min} \\ v_0 \times 10 + \frac{a*10^2}{2} + v_f(t-10); t > 10\mathrm{min}. \end{cases} \tag{6}$$

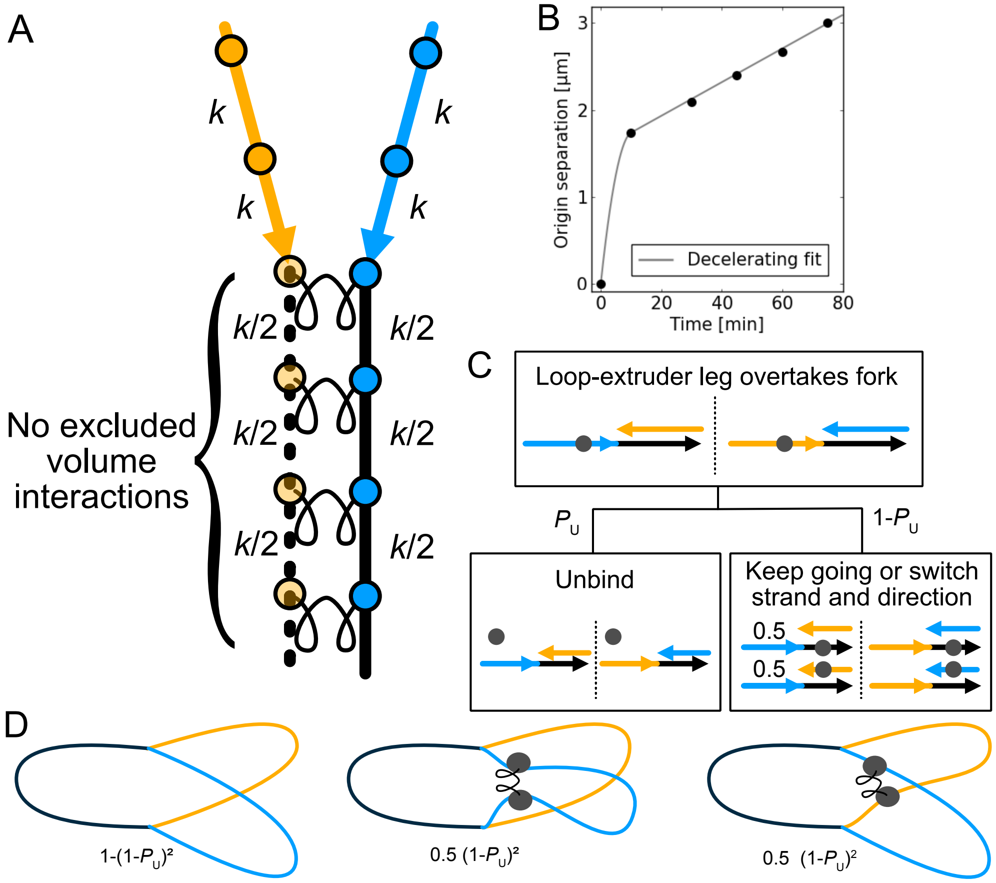


Figure 2: **Simulation details A** The 3D simulations include $2N$ monomers at all time points, some of which may be unreplicated. Unreplicated monomer copies (pale orange) have no excluded volume interactions, and they are tied with springs to their replicated copies (blue). The polymer spring stiffnesses between unreplicated monomers are halved, to account for the fact that there are effectively two springs present in parallel. **B** For simulations where ParAB*S*-like forces were applied to the origins of replication, we assumed initial fast separation of the origins, and afterwards a separation at a constant rate, qualitatively similar to experimental data from [12]. The exact parameters were found as a fit to experimental data [15]. **C** Loop-extruder legs overtaking a replication fork were modeled as either unbinding or jumping onto either of the other two strands at the fork. If the jump is from one replicated strand to the other, the direction of the leg's movement is now from *ter* to *ori*. **D** We consider a loop-extruder loaded at the origin of replication, some time after replication forks have over-taken both of its legs. The loop-extruder can either unbind, have both of its legs jump on the same replicated strand, or have its legs jump on different replicated strands, causing inter-chromosomal contacts.

#### 3.3 Replication speed

We assume that *C. crescentus* replication forks move at equal, constant speeds. Hence, for a genome of size 4.05 Mb, if replication is completed in roughly 75 minutes [11], we expect replication forks to move at speeds of roughly 27 kb/minute. Although replication speeds vary in different growth conditions, we do not expect this parameter to greatly influence our results.

### 4 Description of loop-extrusion simulations for a replicating bacterial chromosome

We edited previously published code for loop-extruders capable of by-passing each other on a bead-spring polymer [7], as follows:

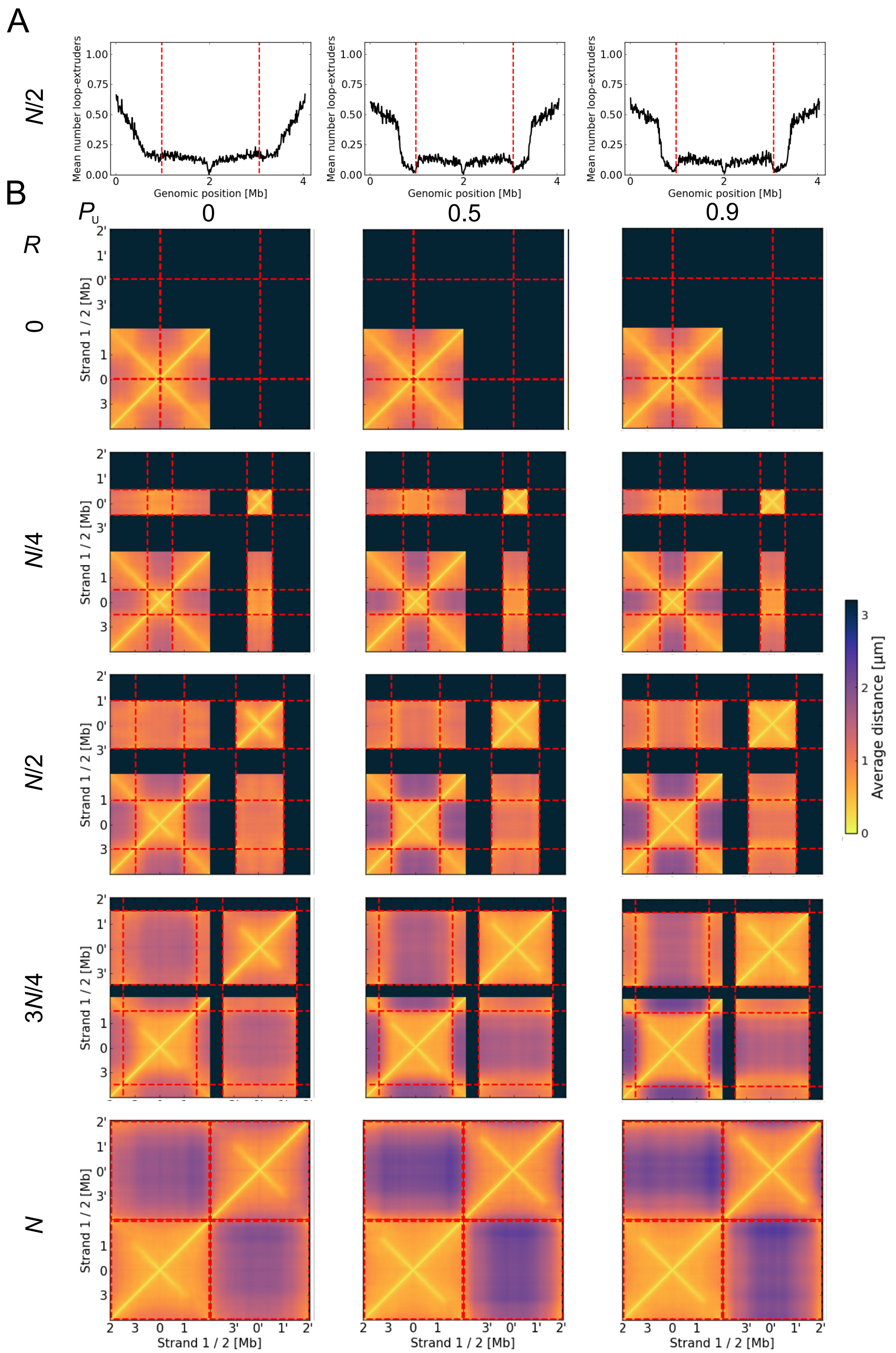


Figure 3: **Unbinding rate at replication forks does not qualitatively affect chromosome dynamics A** A sweep over the unbinding rate of loop-extruders upon replication fork encounter shows that increasing $P_{\mathrm{U}}$ diminishes the number of loop-extruders behind the forks. **B** Distance maps show that the 3D organization of chromosomes is not significantly affected by $P_{\mathrm{U}}$. Larger distances between the replicated strands are observed for higher $P_{\mathrm{U}}$, indicating that segregation is slightly improved.

1. Modelling *C. crescentus*, a single loading site for a fixed number $n_{\mathrm{LE}}$ of loop-extruders at the origin of replication is used. Following [7], on an unreplicated chromosome, loop-extruders have roughly a 50% chance of being loaded at the *ori*, and 50% chance to be loaded elsewhere on the chromosome. The loop-extruder loading affinities of all sites are assumed to stay constant during the replication cycle. Once a monomer is replicated, its new copy is assigned the same loop-extruder loading affinity as the original.
2. The number of monomers corresponding to a single chromosome, $N = 404$, and the average length of a monomer, $b = 129$ nm, are set to describe the mean extension of 10 kb segments in *C. crescentus* [8]. Since plectonemes occur at scales smaller than 10 kb, we do not include plectonemes in our simulations. 1D simulations were performed with 4040 lattice sites for loop-extruders, to ensure that the loop-extruder density and frequency of interactions was the same as in original simulations [7].
3. Each simulation is started with two copies of a chromosome, such that the indices $i$ and $i + N$ reflect an original monomer and a its yet-to-be-replicated copy (Fig. 2A).
4. The loading rates of loop-extruders at unreplicated sites $i + N$ are set to zero.
5. The excluded volume interactions of unreplicated monomers $i + N$ are set to zero.
6. Monomers $i$ and $i + N$ are bound together with a spring of spring constant $k$.
7. Neighboring monomers $i$ and $i - 1$ are bound together by a spring with spring constant $k/2$, as were $i + N$ and $i + N - 1$, such that these parallel springs have a combined strength of $k$ (Fig. 2A).
8. We define replication fork positions $F_1$ and $F_2$, each having a replication speed corresponding to $v_{\mathrm{Rep}} = 1.6 \times v_{\mathrm{LE}}$, where $v_{\mathrm{LE}}$ is the loop-extrusion speed. The ratio comes from dividing the the approximated replication fork speed of 27 kb/min by the rate of condensin zip-up on *C. crescentus* chromosomal arms (17 kb/min, [16]).
9. When a replication fork passes position $i$, we remove the spring between $i$ and $i + N$, turn on excluded volume interactions for monomer $i + N$, and increase the strength of the springs between ($i$,$i - 1$) and ($i + N$,$i + N - 1$) to be of strength $k$.
10. Since the number of loop-extruders increases throughout the cell cycle [7], we assume that the number of loop-extruders per kb stays constant. Hence whenever replication had progressed by an amount $N/n_{\mathrm{LE}}$, we increase the number of loop-extruders by one. At the end of a simulation, the number of loop-extruders has hence doubled.
11. When a replication fork overtakes a loop-extruder leg, or collides with it head-on, the loop-extruder either unbinds with a probability $P_{\mathrm{U}}$ or moves to either the original or replicated copy of the chromosome with equal probabilities (Main Fig.1E).
12. When a loop-extruder leg over-takes a replication fork, it either unbinds with probability $P_{\mathrm{U}}$ or moves to either the unreplicated chromsome strand or the other replicated chromosome strand, with equal probabilities (Fig. 2C). Note that if the other replicated strand is chosen, the direction of travel of the loop-extruder leg has to be switched, since the newly replicated chromosomal strands converge at the replication fork.
13. In simulations with origin-pulling forces, we add forces that hold the origins of replication at given distances with respect to the cell pole. See Section 2 6 for details.

We note that our chosen behavior for what happens during encounters between replication forks and loop-extruders rests on three assumptions: loop-extrusion is non-topological (See Section 12 for a topological alternative); loop-extruder legs can by-pass replication forks with some probability; and that after by-passing a fork, a loop-extruder leg cannot distinguish between two chromosomal strands it encounters. This scheme results in a fraction of loop-extruders overtaken by the replication forks connecting newly replicated sister chromosomes, hence inhibiting chromosome segregation (Fig. 2D). This effect diminishes as $P_{\mathrm{U}} \to 1$ or if the loop-extruder leg and replication fork speeds are similar (in this case, an over-taken loop-extruder "keeps up" with the replication forks).

Given these considerations, simulations were run as follows:

1. Initialize positions of two tied-together copies of the chromosome in a cylindrical confinement and in an *ori-ter* configuration.
2. Perform loop extrusion simulations without replication for more than $N/2/v_{\mathrm{LE}}$ simulation steps; long enough for loop-extruders to move all the way across the chromosome.
3. Start movement of replication forks.
4. Save the positions of loop-extruders and the replication forks every few simulation time-steps.

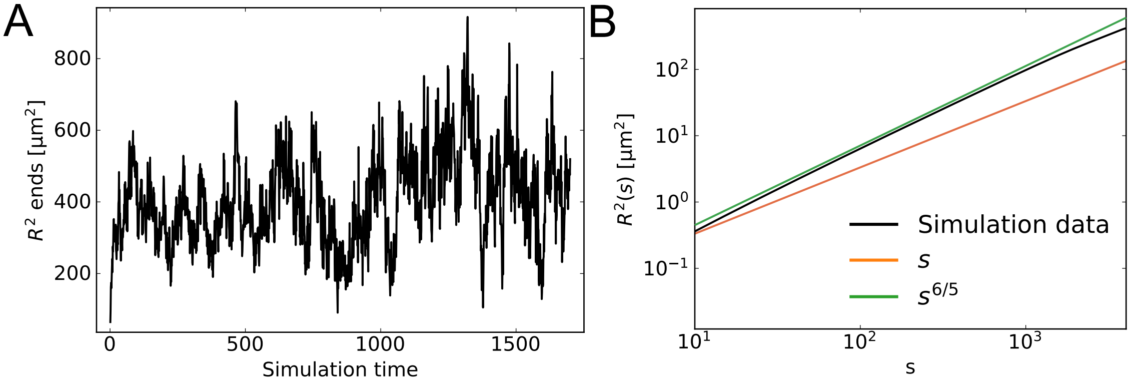


Figure 4: **Mean-squared distance scaling A** The convergence of the end-to-end mean-squared distance for a linear polymer of length $N = 4040$, without confinement. The configuration was initialized as a random walk. **B** The converged (simulation time$> 500$) curve for the $R^2(s)$ of the unconfined linear polymer. Colored lines show the expected scaling for an ideal chain ($s$) and a self-avoiding chain ($s^{6/5}$). For large $s$, the simulations converge to the self-avoiding limit.

5. Run simulations until replication is complete, or for around $N/2/v_{\mathrm{Rep}}$ time-steps.
6. Start 3D simulations.
7. For each simulation step, fetch saved positions of loop-extruders and replication forks at a saved time-point.
8. Add springs between monomers tied together by a loop-extruder.
9. Given a list of monomers replicated since the last step, turn on their excluded volume interactions and edit the spring strengths as described in Fig. 2A.
10. Given the current replication stage, calculate the current size of the cell (See Section 3 for details).
11. Given the new cell size, increase the confinement length at both ends of the cell symmetrically.
12. If using origin-pulling forces, infer the new distance between the *ori*s, update the tether forces that hold them at the appropriate heights.
13. Perform 3D polymer simulation steps.
14. Save the polymer configuration alongside the current positions of the loop-extruders and replication forks.

The code used in the simulations can be found at github.com/PLSysGitHub/loop-extrusion_with_replication.

### 5 Sufficiency of excluded volume interactions

To ensure that the excluded volume interactions are strong enough to give self-avoiding behavior for the chain, we simulated a linear polymer of length $N = 4040$ without confinement, and analyzed the mean-squared distance between monomers a genomic distance $s$ apart:

$$R^2(s) = \left\langle (\mathbf{x_i} - \mathbf{x_{i+s}})^2 \right\rangle \quad (7)$$

where the average is taken over all monomers $i$ and multiple converged steady state simulations (Sup. Fig. 4A).

For sufficiently large $s$, ideal chains with no excluded volume interactions, $R^2 \propto s$. For self-avoiding chains, on the other hand, $R^2 \propto s^{2\nu}$, with $\nu \approx 3/5$. The excluded volume interactions in our simulations show self-avoiding scaling for large $s$ (Sup. Fig. 4B), and hence we conclude that the excluded volume interactions are strong enough to give self-avoiding chain behavior. Note, the excluded volume interactions are kept low enough so that monomers can occasionally move through each other [6]. Such an approach is justified at the coarse-graining level of the simulations: While the polymers cannot overlap at the molecular scale, physically the course-grained representation of the strands are allowed to overlap in space, but with an energy penalty set by the excluded volume interaction.

### 6 Origin-pulling forces

For simulations with origin-pulling, we needed to constrain the expected long axis separation between origin replicates, $d_{\mathrm{ori}}$ as a function of replication stage (Section 3.2). Origin 1 was then constrained by a harmonic force that pulled it

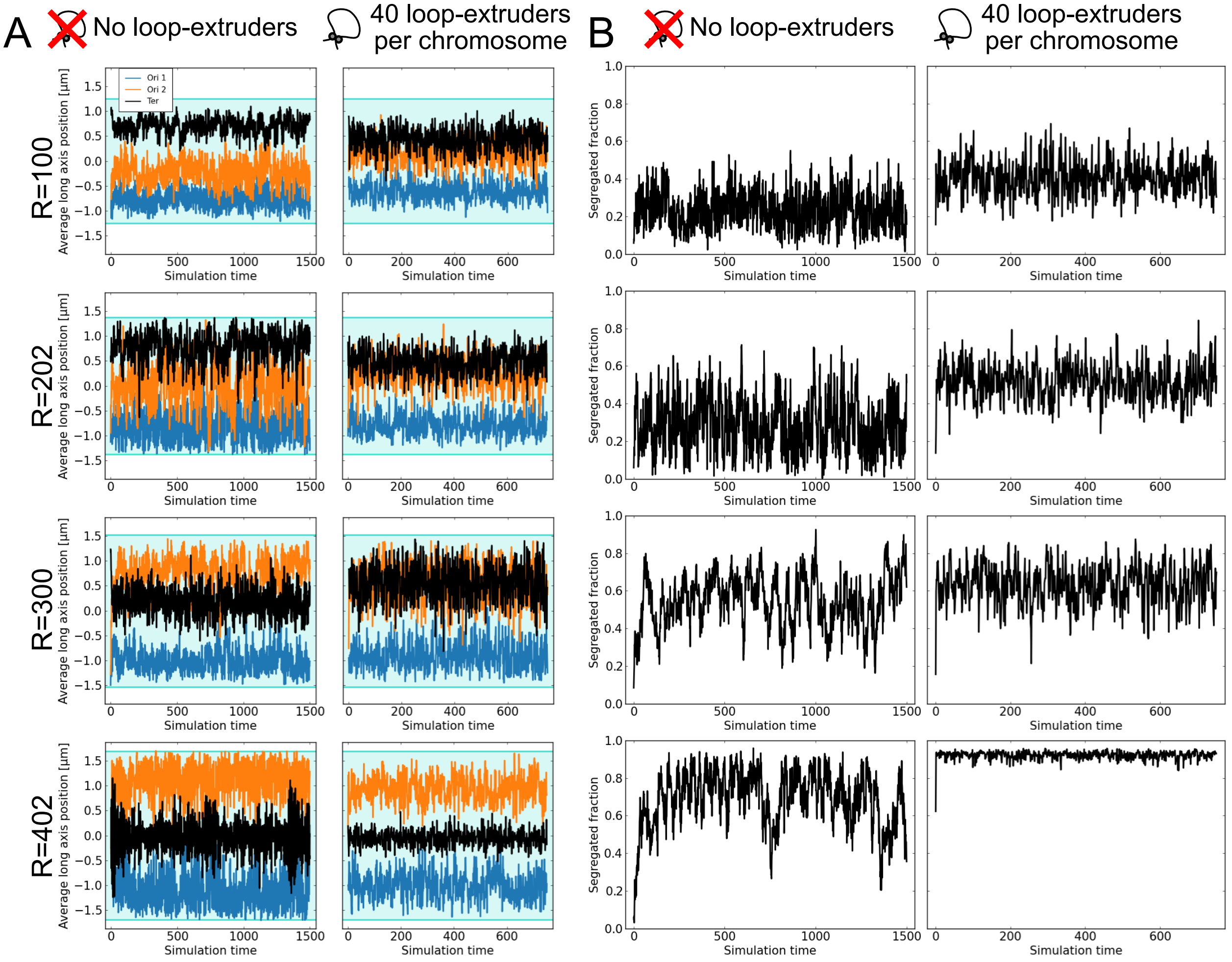


Figure 5: **Convergence of steady-state simulations A** Tracks of the mean long axis positions of monomers over simulation time for steady state simulations. All chromosomes were initialized in an unsegregated *ori*-*ter* configuration. For most replication stages $R$, convergence was reached around 100 simulation steps. For $R = 402$ without loop-extruders, fluctuations were higher, and convergence was only reached after approximately 300 time steps. **B** The fraction of segregated monomers (see Main Figure 2G) was also found to converge after around 100 simulation steps.

towards a height 343 μm from Pole 1 [8], whereas Origin 2 was constrained by a harmonic force that pulled it towards a height $343 + d_{\text{ori}}$ μm from Pole 1.

### 7 Predicting fork separation

Consider two replication forks separated by a linear segment in the absence of loop-extruders. If the long-axis extension per monomer is constant, we expect that a replicated strand of length $R < N/2$ should extend by $L_{\text{max}} 2R/N$, and an unreplicated strand of length $N - R < N/2$ by $L_{\text{max}}(N - R)/R$, where $L_{\text{max}}$ is the expected distance between two monomers at opposite poles of the cell. Here we use a maximum distance of $L_{\text{max}} = L - d$, since each monomer in on average in the middle of a confinement blob of radius $d/2$, and hence on average at least a distance $d/2$ from the cell pole. Similarly, in the presence of loop-extruders, we expect that for $R < N/2$, the *ori*-fork separation is $L_{\mathbf{max}} R/N$. Once $R \geq N/2$, the chromosome adopts an *ori*-*ter*-*ori* configuration, and the replication forks should be located mid-cell. We note that this is a simple approximation; a proper analysis should account for excluded volume interactions and the effect of boundaries on the monomer density.

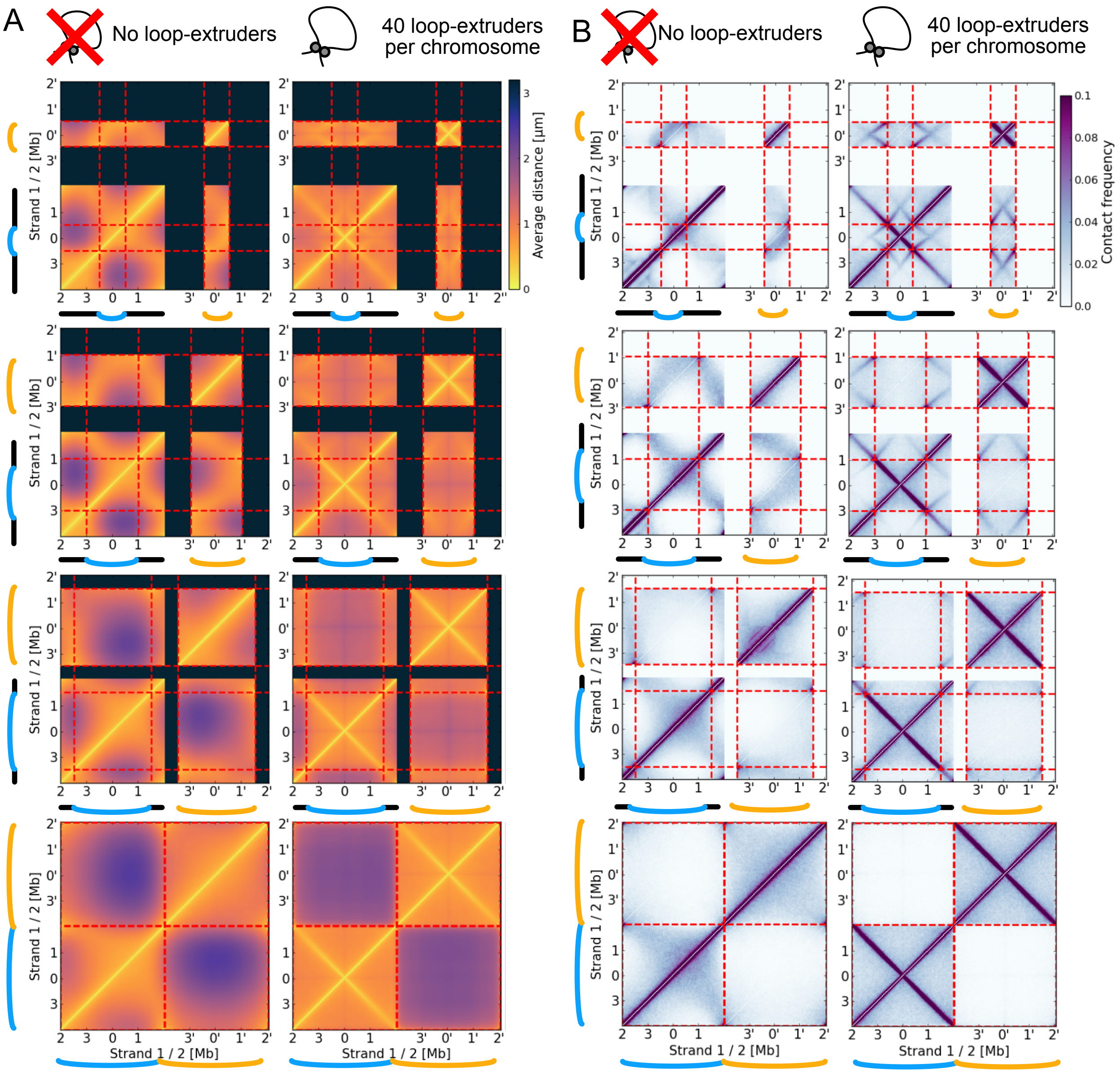


Figure 6: **Distance and contact maps from steady-state simulations A** Distance maps from steady-state simulations without and with loop-extruders. The value at a point $(i, j)$ gives the average 3D distance between monomers $i$ and $j$. Black sections on the map represent unreplicated chromosome segments, dashed red lines give replication fork positions. All analyzed configurations were oriented as described in Section 7. **B** Steady-state simulation data was used to calculate Hi-C-like contact maps without or with loop-extruders. A "contact" was defined as an event where two monomers were within a two monomer length distance of each other (258 nm). Main Figure 2D shows zoom-ins of the inter-chromosomal contacts, here the top-left squares of the full contact maps.

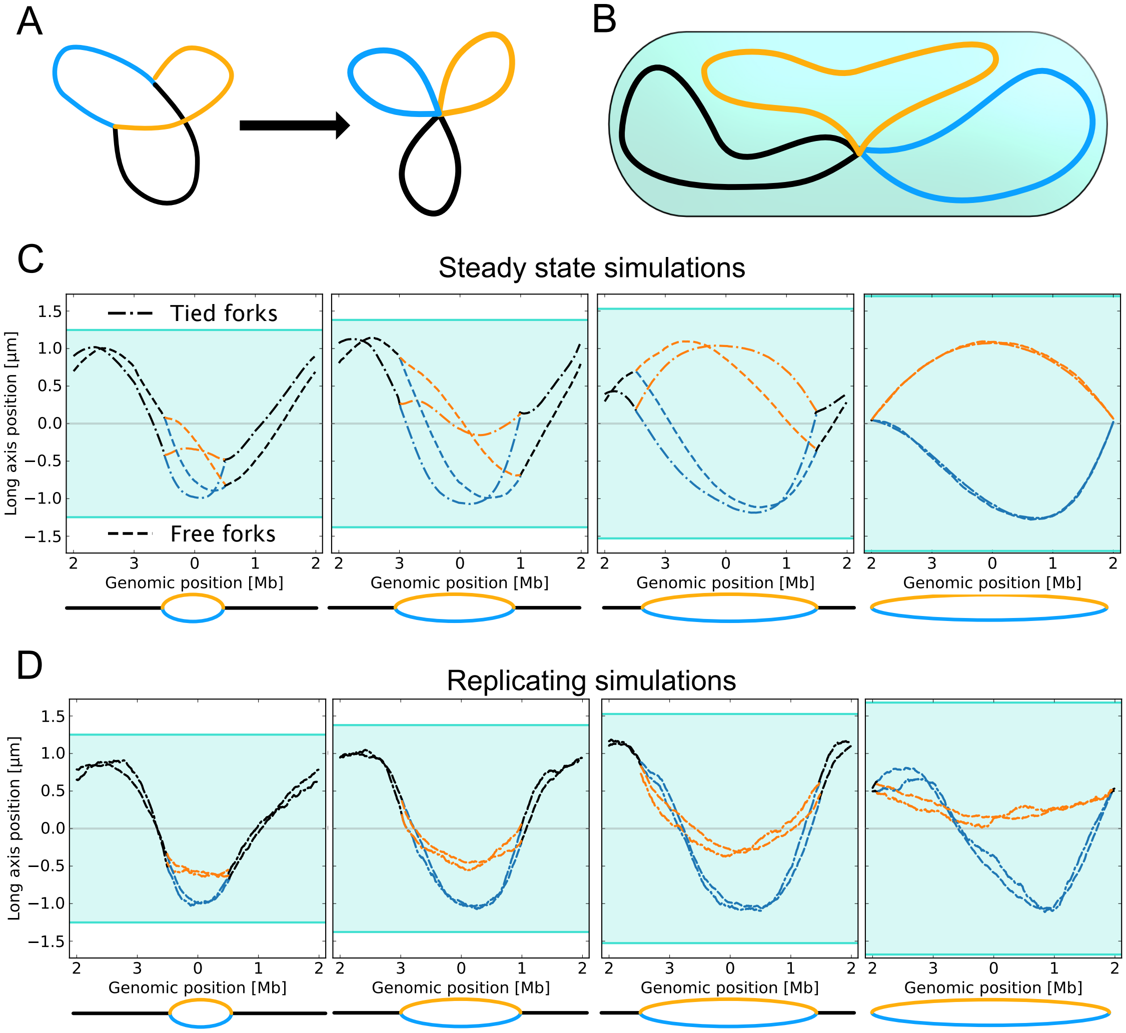


Figure 7: **Replication factory model A** The replication factory model was created by connecting the replication forks of the polymer model with a spring. No loop-extruders were used in the simulations. **B** A sketch of the converged configuration in the replication factory model, for $R = N/2$. **C** Mean long axis positions of monomers from simulations with no loop-extruders, with or without a replication factory. **D** Mean long axis positions of monomers from dynamically replicating simulations with no loop-extruders, with or without a replication factory.

### 8 Replication factory simulations

In addition to linearizing the origin-proximal regions of replicating chromosomes, loop-extruders loaded at the origins of replication will confine replication forks to be spatially proximate. Our loop-extrusion model hence gives rise to an effective "replication factory"; the two replisomes are spatially close to each other, as seen in some bacteria [17, 18]. Such constraints on the distance between replication forks have been simulated before, but they were insufficient to give concurrent replication and segregation without an additional concentric-shell mechanism [19]. In another simulation study, confining replication forks to mid-cell and rescaling simulation time allowed for entropic segregation concurrent with replication [20]. We note, however, that this study used a much lower monomer count of 80.

To explore the direction of entropic forces in the replication factory model, where entropically preferred fork segregation is not possible, we conducted both steady-state and replicating simulations without loop-extruders, but with the replication forks tied together by a spring (Sup. Fig. 7A).

In the steady-state simulations, we found that including a replication factory led to better segregation in the absence of loop-extruders (Sup. Fig. 7C). At the final replication stages, even without a replication factory, the forks are close to each other at the terminus, and the simulations show similar configurations. Interestingly, half-way through replication,

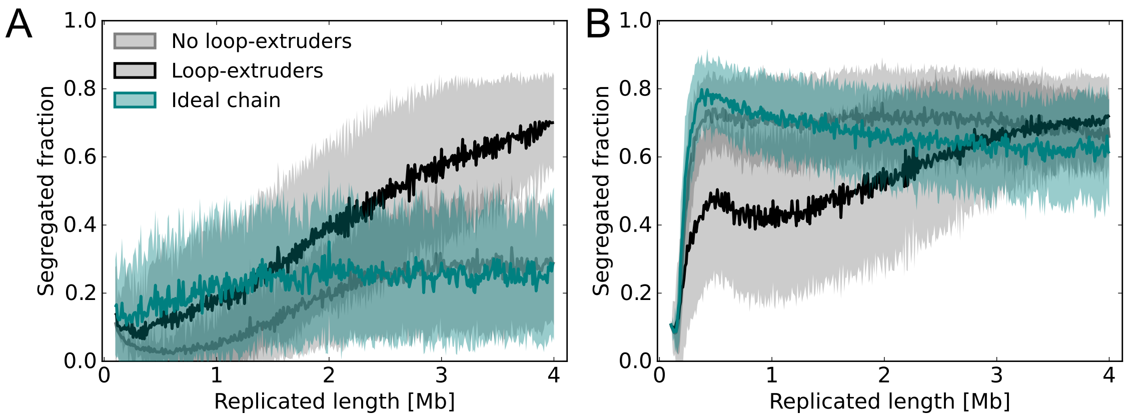


Figure 8: **Standard deviations of ideal chain segregated fractions A** Subfigure 2C, showing the segregated fraction for replicating simulations without *ori*-pulling, including the standard deviation for ideal polymer simulations. **B** Similarly for Subfigure 2F; the segregated fraction for replicating simulations with *ori*-pulling.

the entropically preferred configuration showed one newly replicated DNA strand spread out symmetrically around mid-cell (Sup. Fig. 7B). Inclusion of loop-extruders prevents this spreading out, since loop-extruders tie the arms of the newly replicated strand together.

In replicating simulations, including a replication factory did not noticeably improve segregation (Sup. Fig. 7D). The newly replicated strands were spread around mid-cell, and as a result did not have time to segregate even after replication proceeded further. This suggests that loop-extrusion mainly aids in segregation by linearizing origin-proximal regions, rather than by tying the replication forks together.

### 9 Time-scale of simulations and faster growth

To assess how the time-scales of our polymer simulations compare to the relaxation times of actual bacterial chromosomes, we analyzed the mean-squared displacement (MSD) $\langle |\mathbf{R}(t) - \mathbf{R}(0)|^2 \rangle$ of monomers in the first minutes of replicating simulations as a function of time (Fig. 9A,B). At these time-scales, replication and cell growth do not significantly affect the monomer dynamics, and hence we expect that monomers diffuse according to a power-law:

$$\langle |\mathbf{R}(t) - \mathbf{R}(0)|^2 \rangle = 4Dt^{\gamma}, \tag{8}$$

where $D$ is the apparent diffusion constant, and $\gamma$ is an exponent determined by whether the monomer motion is diffusive $(\gamma = 1)$ or sub-diffusive $(\gamma < 1)$.

We compared our simulations to experimental MSD data for labels placed near the origin of replication in *E. coli* cells [21]. With loop-extruders but no *ori*-pulling forces, our simulations showed an *ori* diffusivity close to experiment, but we found that monomer diffusion coefficients varied in a broad range of $10^{-3}$ to $10^{-2}$ $\mu\text{m}^2\text{s}^{-\gamma}$ (Fig. 9C). The fitted exponents $\gamma$ mostly corresponded sub-diffusive motion, with values in the range $0.2 - 0.7$ (Fig. 9D), with the peak of the distribution around the experimentally observed $0.39 \pm 0.04$ [22].

An average diffusion coefficient of order $0.01$ $\mu\text{m}^2\text{s}^{-\gamma}$ would give a time scale of minutes for entropic segregation, as estimated by [19]. We hence expect entropic forces to act fast enough at the scale of the simulations, which explains why the replicating simulations show transitions towards fork-segregated states in the absence of loop-extruders.

To see if our findings were dependent on the time-scales of our simulations, we slowed down the 3D polymer dynamics by increasing excluded volume interactions, by decreasing the number of 3D polymer steps per 1D loop-extrusion step, as well as by increasing the drag coefficient in our simulations( Fig. 9A). We found that in the presence of loop-extruders, chromosome segregation remained effective in all cases, but in the absence of loop-extruders, segregation was inhibited (Fig. 9D,E).

### 10 Number of loop-extruders

It is experimentally difficult to estimate the density of loop-extruding condensins on a bacterial chromosome. We used a density of 40 loop-extruders per chromosome, an estimate for *B. subtilis* based on matching simulations to

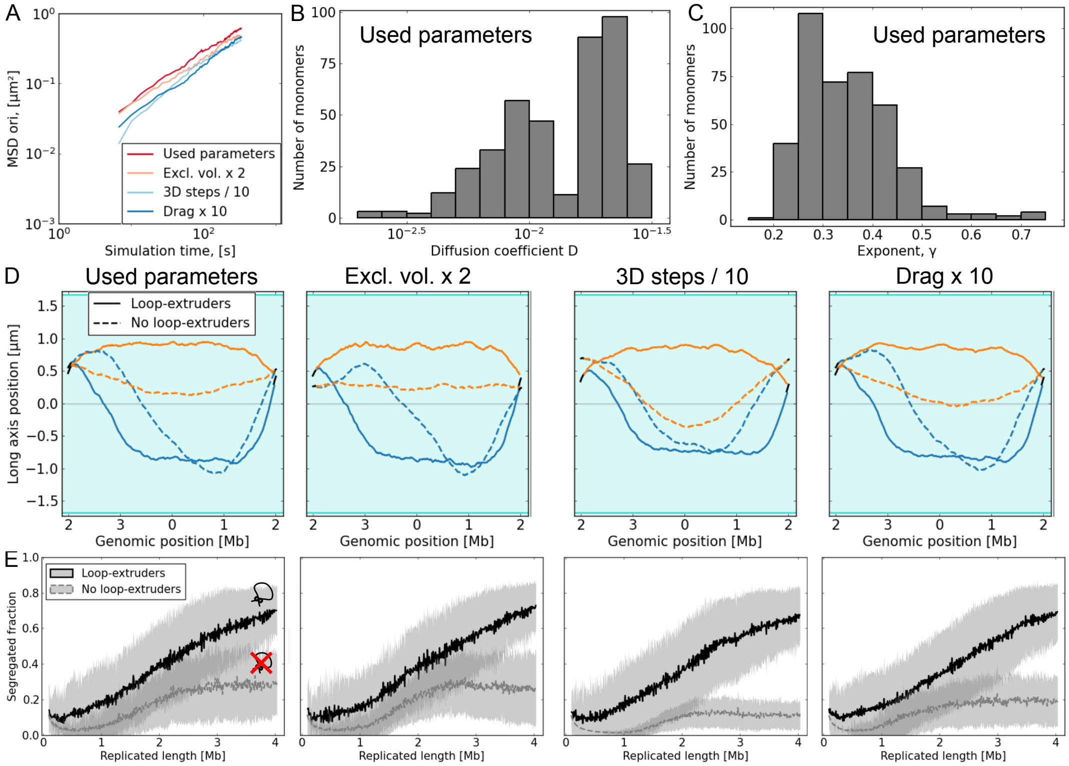


Figure 9: **Time-scales of simulations A** The mean-squared displacement (MSD) for the origin as a function of time. The MSD is of the same order of magnitude as found by [21]. By varying simulation parameters, we were able to slow down the polymer dynamics relative to loop-extrusion, or equivalently, to consider faster replication. **B** The distribution of diffusion coefficients $D$ for the parameters used in most simulations. The total number of monomers is 404. **C** The distribution of exponents $\gamma$ for the parameters used in most simulations. **D** When polymer dynamics were slowed down in replicating simulations, the terminal regions of chromosomes appeared more intermingled in the absence of loop-extruders. In the presence of loop-extruders, however, no qualitative difference was found. **E** In replicating simulations where the number of 3D steps per minute was reduced, or the drag coefficient was increased, the segregated fractions of monomers slighly decreased in the absence of loop-extruders. However, in the presence of loop-extruders, segregation remained efficient.

experimental Hi-C data [6, 7]. However, it is possible that the loop-extruder density in *C. crescentus* differs from this value. We therefore repeated our replicating simulations with half and twice as many loop-extruders. The over-all organization of the chromosomes did not change significantly over time, and the replicated fractions were also found to be similar (Fig. 10).

### 11 Decreasing the range of loop-extruder progression along the chromosome

In *C. crescentus*, condensins appear to typically unbind before progressing up to a characteristic distance of roughly 600 kb from the origin of replication [16]. The blob argument in Section 1 suggests that even partial linearization of the replicating chromosome will make segregated chromosome configurations favorable compared to fork-segregated ones. To test whether such partially linearized chromosomes could still segregate, we simulate replication processes with different life-times $\tau$ for loop-extruders (Sup. Fig. 11). This means that loop-extruders travel a shorter average distance $\lambda \approx v_{\text{LE}}\tau$ before unbinding from the DNA. We find that even $\lambda/N = 0.15$ (close to $600/4040$ for *C. crescentus*) is sufficient for segregation.

### 12 Topological loop-extruders

It is currently unknown what the exact mechanism for condensin loop-extrusion is. Some have hypothesized that a condensin leg forms a ring around DNA, and that this ring is not opened during loop-extrusion, resulting in so-called

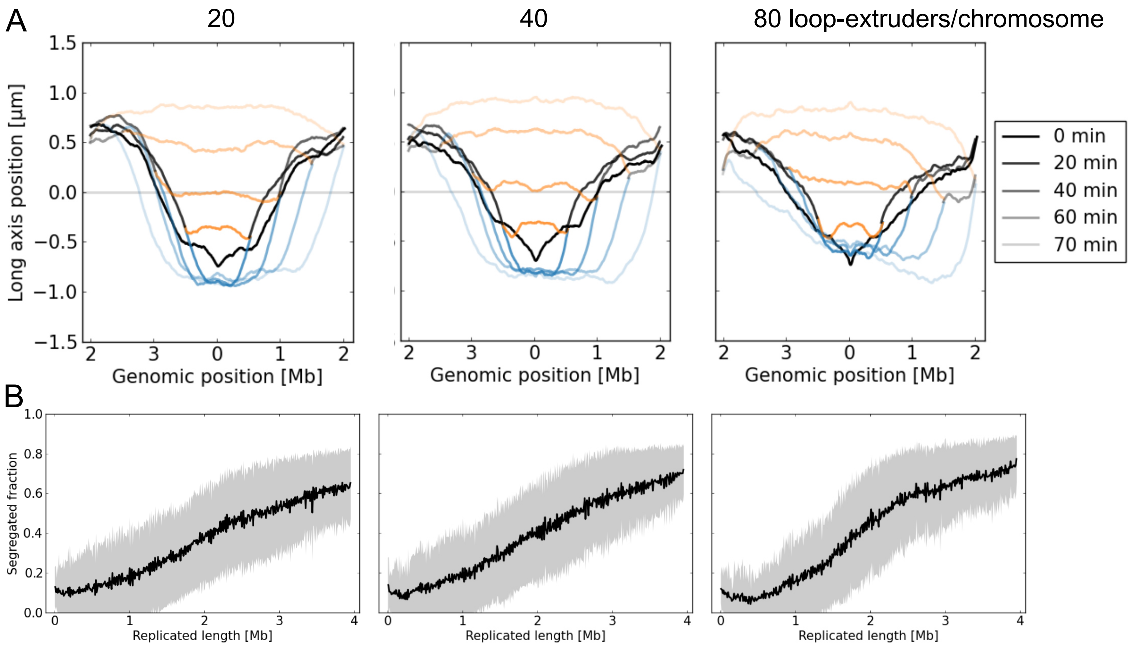


Figure 10: **Number of loop-extruders does not significantly affect chromosome dynamics A** Increasing or decreasing the density of loop-extruders in the replicating simulations does not have a qualitative impact on the long-axis organization the of chromosomes. However, a smaller number of loop-extruders appears to give slightly better segregation, as seen by the increased separation between origins of replication. **B** Changing the number of loop-extruders does not significantly affect the segregated fraction over the replication cycle.

"topological loop-extrusion". This model contradicts evidence that yeast condensin is able to slide over very large obstacles [23] and both bacterial and eukaryotic are capable of by-passing each other [7, 24]. We nevertheless considered an alternative simulation scheme with topological loop-extruders; loop-extruders are not allowed to by-pass each other, and when a loop-extruder encounters a replication fork, it encircles two strands of DNA on the other side (Fig. 13A, B).

We note that such loop-extruder legs contain only one or two DNA strands as long as loop-extrusion is fairly symmetric. To see this, we consider what happens when a loop-extruder takes over a fork from either a replicated or an unreplicated strand.

First, suppose that a loop-extruder leg approaches a fork from the unreplicated side of the chromosome. It then encloses two newly replicated segments. If it traverses these two *equal length* segments at the same speed, upon reaching the next replication fork, it will simply enclose a single unreplicated strand again (Fig. 13C).

Now suppose that a loop-extruder is loaded behind a replication fork at $F_1$, such that one leg immediately overtakes the fork. Since the other leg of the loop-extruder must travel a distance less than or equal to $R$ to reach $F_2$, the two legs collide before the first leg can engulf another strand of DNA (Fig. 13D). Since topological loop-extruder legs cannot traverse one another, a third strand is thus never engulfed.

Motivated by this finding, we edit the loop-extrusion simulations as follows:

1. The by-passing probability for loop-extruders is set to zero.
2. Instead of having vectors of dimension one to track the positions and directions of loop-extruder legs, we now track two positions and two directions per chromosomal leg. The two values are set to be the same whenever a loop-extruder leg only encompasses one chromosomal strand.
3. When a fork overtakes/collides head-on with a loop-extruder at position $i$ with only one unreplicated strand in it, we add the newly replicated monomer $i + N$ to the leg (Fig. 13A). The direction of travel on both strands remains the same.
4. When a loop-extruder with one replicated strand in it overtakes the fork at $F_1$ (or $F_2$), we add the strand $F_1 + N - 1$ (or $F_2 + N + 1$) to the loop-extruder leg (Fig. 13B). The direction of travel on this second strand has to be switched.

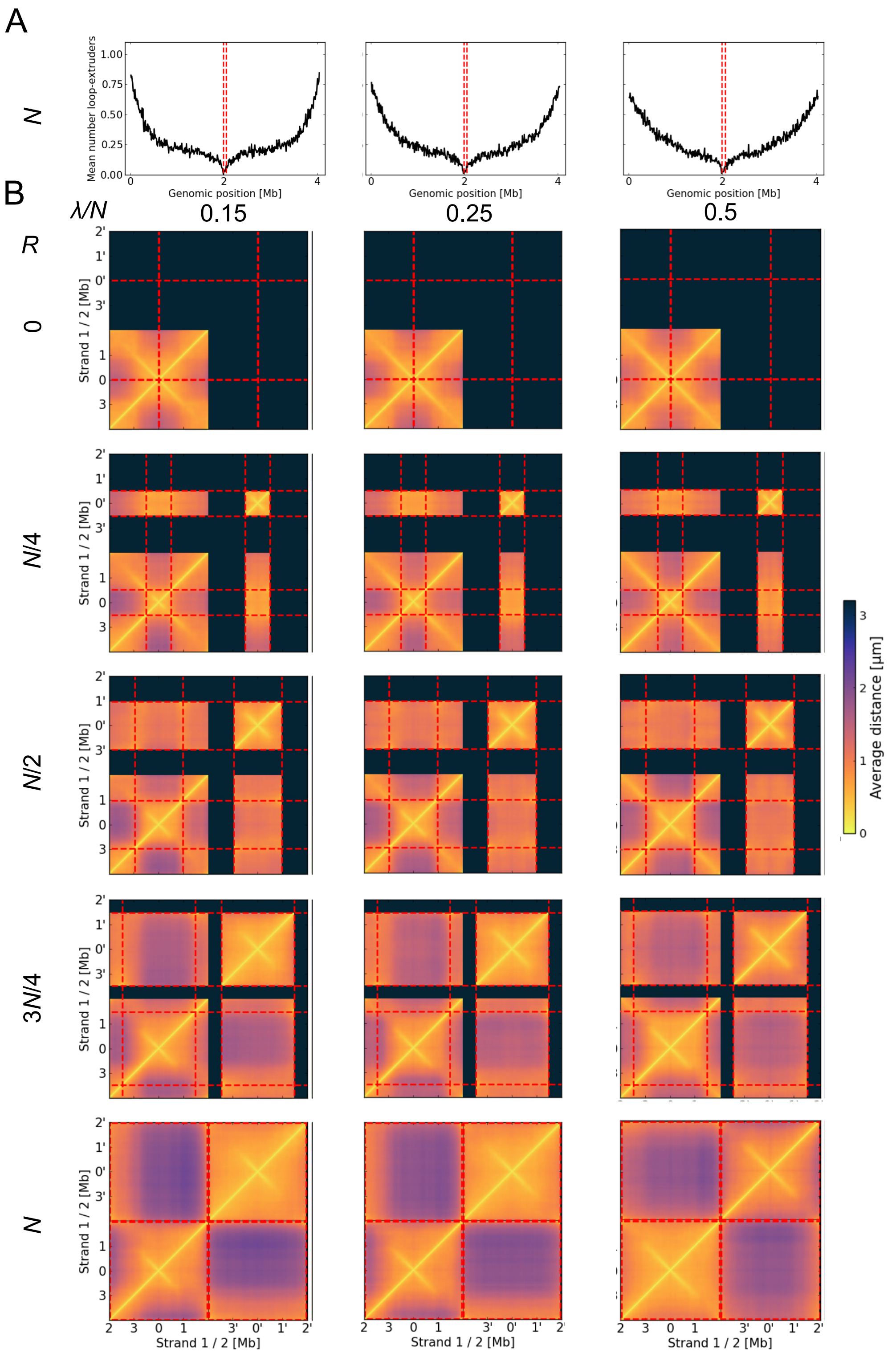


Figure 11: **Faster loop-extruder unbinding still allows for concurrent replication and segregation A** The average number of loop-extruders at different genomic positions when $R = N$, for different loop-extruder lifetimes. **B** Distance maps for replicating simulations with different loop-extruder lifetimes.

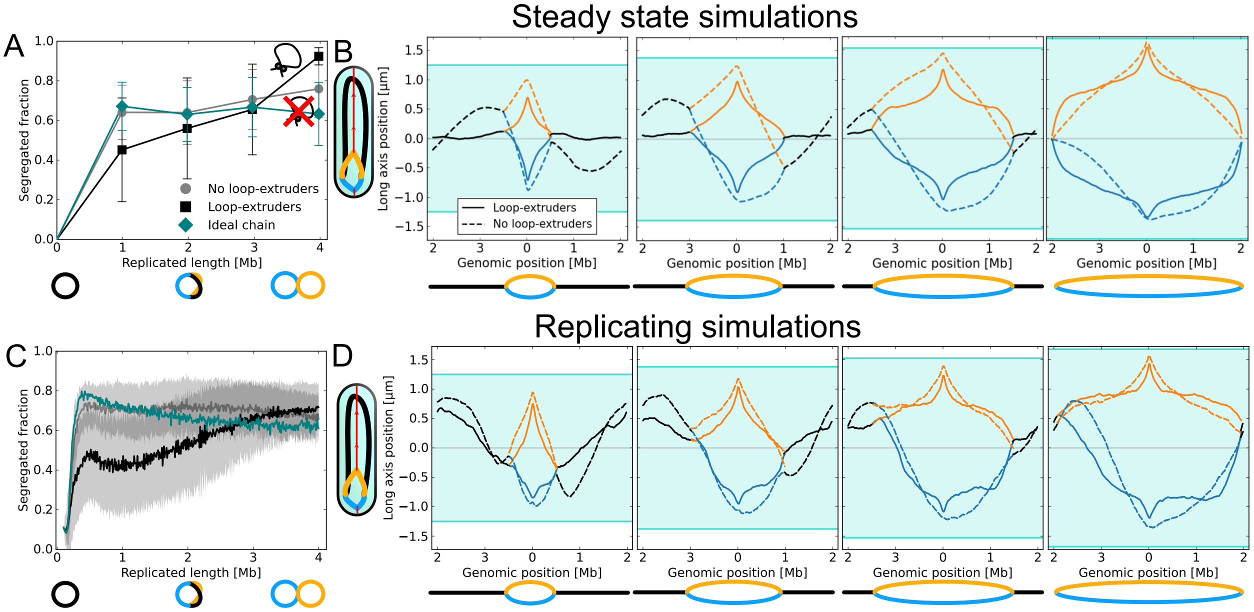


Figure 12: **Simulations with *ori*-pulling forces A** Similar to Main Figure 2G; in the presence of origin segregation forces, steady state chromosomes are more strongly segregated, especially at early replication times. However, near the end of replication, origin segregation forces no longer seem relevant. **B** Similar to Main Figure 2A. Mean long axis positions of loci reveal that although origin-pulling forces help segregate origin-proximal regions, without loop-extruders we still see significant fork segregation. **C** Similar to Main Figure 3C. In replicating simulations, the segregated fraction increases faster without loop-extruders, but starts to decrease at later replication stages. See Sup. Fig. 8 for a figure with the standard deviation for the ideal chain. **D** Similar to Main Figure 3D. The mean long axis positions with *ori*-pulling, with or without loop-extruders.

5. When a loop-extruder with two replicated strands in it overtakes a fork $F_i$, we set both strands in the loop-extruder to $F_i$.
6. Otherwise the movements of loop-extruders with a single strand in them remain the same.

Since for *C. crescentus* condensin seems to extrude loops at a rate slower than the replication fork speed, replication forks are mostly over-taking loop-extruders or colliding with them head-on. In such a case, topological loop-extrusion is expected to enhance contacts between the two replicated strands of a chromosome, thus inhibiting chromosome segregation. Furthermore, when loop-extruders are not able to by-pass each other, traffic jams will arise near the replication forks, where condensins loaded at two different origins will collide.

In *B. subtilis*, condensins move at speeds of approximately 52 kb per minute [25], faster or comparable to the speed of the replication forks [26]. If the speed of a topological loop-extruder would be higher than that of a replication fork, a loop-extruder loaded at an *ori* after the start of replication would eventually tie together a newly replicated copy of a chromosome with the unreplicated part. Unless loop-extruders would be loaded at a single *ori*, this would also inhibit chromosome segregation.

We hence conclude that irrespective of loop-extrusion speed, topological loop-extrusion at the replication forks is expected to inhibit chromosome segregation in bacteria. This effect is diminished if the loop-extruders unbind at the replication forks, or have similar speeds as the replication forks, although head-on collisions between replication forks and loop-extruders would still result in effective links between newly replicated chromosomes.

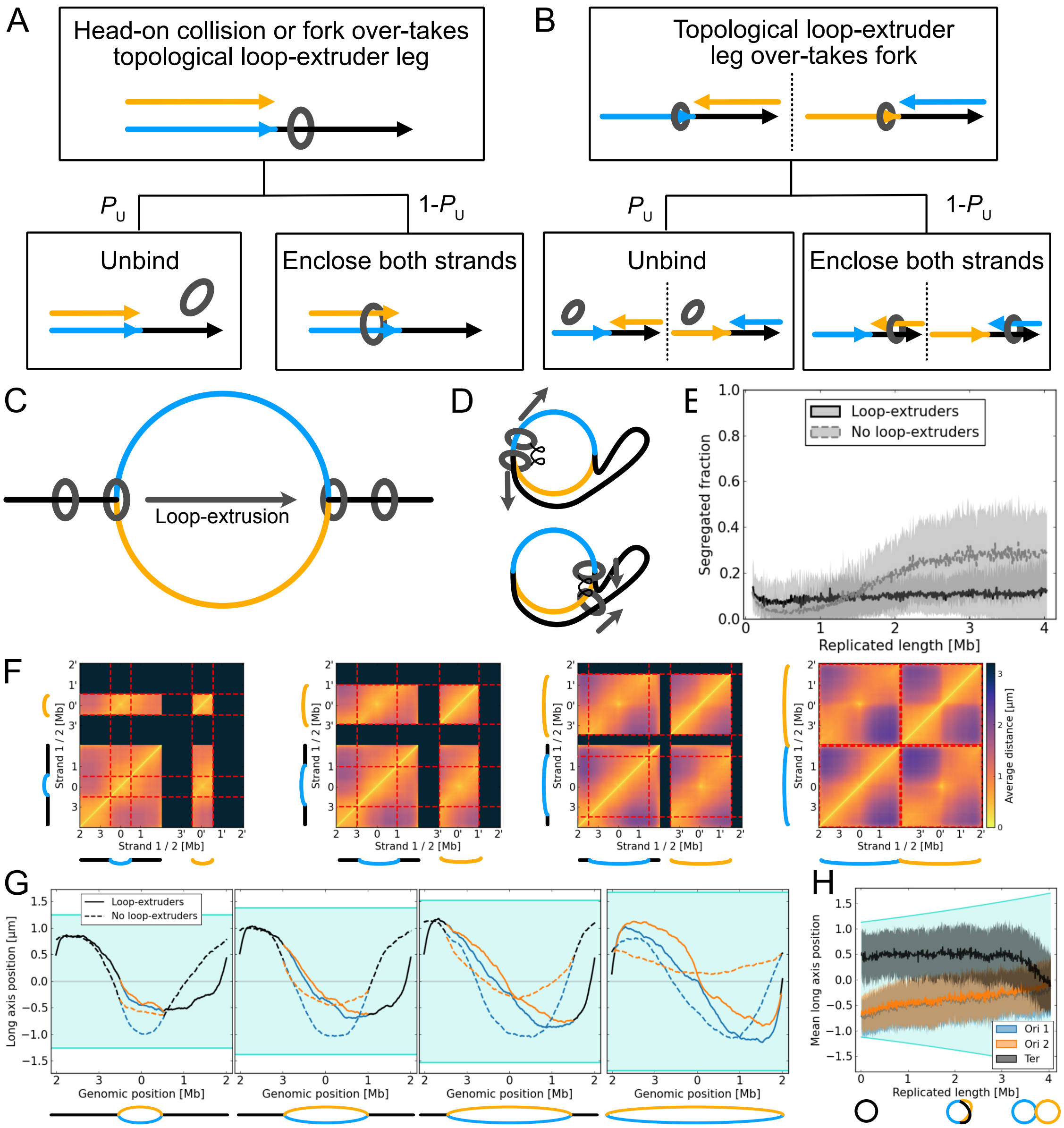


Figure 13: **Details and results from simulations of topological loop-extrusion A** When a replication fork over-takes a topological loop-extruder leg, both the loop-extruder either falls off, or engulfs both newly replicated strands. **B** When a topological loop-extruder leg over-takes a replication fork, it either falls off or engulfs the unreplicated and other replicated strand. **C** A topological loop-extruder leg that collides head-on with a replication fork can only engulf a maximum of two chromosomal strands, since both newly replicated strands are of the same length, so that the loop-extruder will again engulf a single strand at the once it reaches the other fork. **D** A topological loop-extruder that over-takes a replication fork can only engulf a maximum of two strands. Suppose loop-extruder leg 1 over-takes a fork right after loading. If the other leg 2 travels at the same speed, 1 and 2 will collide before 1 reaches the second replication fork. **E** In dynamic simulations, topological loop-extruders were seen to slow-down segregation compared to simulations without loop-extruders. This contrasts the case of nontopological loop-extruders, which significantly sped up segregation (Main Figure 3C). **F** Distance maps from dynamic simulations with topological loop-extruders show that the two replicated chromosomal strands are mostly aligned. In addition, the left and the right arm of the chromosome separate, indicative of a left-*ori*-right organization. **G** The mean long-axis positions of monomers show that, in the presence of topological loop-extruders, the *ter* and both copies of the origin are roughly at mid-cell, whereas the left-and right arms of the chromosomes are in opposite cell halves. **H** Tracks of the origins and terminus in the presence of topological loop-extruders show a transition towards left-*ori*-right configuration, with both *ori*s and the *ter* located roughly at mid-cell.

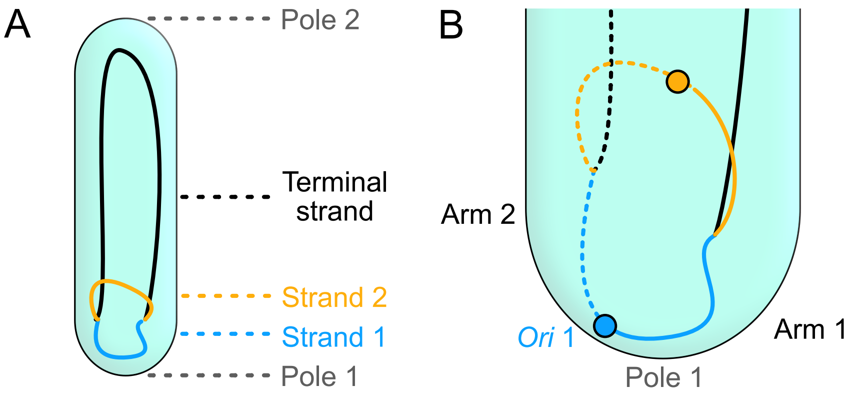


Figure 14: **Relabelling chromosomal strands and arms A** We label Strand 1 as the replicated strand with the center of mass of Strand 1 and the unreplicated strand closer to mid-cell. Pole 1 is the pole closer to Strand 1. **B** After labeling Strand 1 and Strand 2, we label Arm 1 as the arm of Strand 1 and the unreplicated strand that is closer to Pole 1 (full line; dashed line for Strand 2). Note that relabelling of chromosomal arms merely corresponds to changing monomer indices $1, 2, 3..., N$ to run in reverse.

[14] Hoong Chuin Lim, Ivan Vladimirovich Surovtsev, Bruno Gabriel Beltran, Fang Huang, Jörg Bewersdorf, and Christine Jacobs-Wagner. Evidence for a DNA-relay mechanism in ParABS-mediated chromosome segregation. *eLife*, May 2014.

[15] Janni Harju, Joris J. B. Messelink, Grzegorz Gradziuk, Muriel C. F. van Teeseling, and Chase P. Broedersz. In preparation, 2023. Data on separation between origins of replication over the cell cycle.
